# Structural basis of the regulation of normal and oncogenic methylation of nucleosomal histone H3 Lys36 by NSD2

**DOI:** 10.1101/2020.12.05.413278

**Authors:** Ko Sato, Amarjeet Kumar, Keisuke Hamada, Chikako Okada, Asako Oguni, Ayumi Machiyama, Shun Sakuraba, Tomohiro Nishizawa, Osamu Nureki, Hidetoshi Kono, Kazuhiro Ogata, Toru Sengoku

**Affiliations:** Department of Biochemistry, Yokohama City University Graduate School of Medicine; Institute for Quantum Life Science, National Institutes for Quantum and Radiological Science and Technology; Department of Biological Sciences, Graduate School of Science, The University of Tokyo

## Abstract

Dimethylated histone H3 Lys36 (H3K36me2) regulates gene expression by antagonizing the repressive effect of polycomb-group proteins. Aberrant upregulation of H3K36me2, either by overexpression or point mutations of NSD2/MMSET, an H3K36 dimethyltransferase, is found in various cancers, including multiple myeloma. To understand the mechanism underlying its regulation, here we report the cryo-electron microscopy structure of the catalytic fragment of NSD2 bound to the nucleosome at 2.8 Å resolution. The nucleosomal DNA is partially unwrapped at superhelix location +5.5, facilitating the access of NSD2 to H3K36. NSD2 interacts with DNA and H2A along with H3. The autoinhibitory loop of NSD2 changes its conformation upon nucleosome binding to accommodate H3 in its substrate-binding cleft. Kinetic analysis revealed two oncogenic mutations, E1099K and T1150A, to aberrantly activate NSD2 by increasing its catalytic turnover but not the nucleosome affinity. Molecular dynamics simulations suggested that in both mutants, the autoinhibitory loop adopts an open state that can accommodate H3 more often than the wild type. We propose that E1099K and T1150A destabilize the interactions that keep the autoinhibitory loop closed, thereby enhancing the catalytic turnover. Our analyses would guide the development of specific inhibitors of NSD2 for the treatment of various cancers.

## Introduction

Histones are subjected to a variety of post-translational modifications that regulate diverse aspects of genome functions (1). Dysregulation of histone modifications is often linked to diseases, such as developmental defects and cancers (2). NSD2 (also known as WHSC1/MMSET) is a member of the NSD family that catalyzes the mono- and di-methylation of histone H3 K36 (3). Dimethylated H3 K36 (H3K36me2) antagonizes the activity of polycomb repressive complex 2 (PRC2) in catalyzing H3K27 trimethylation, a hallmark of repressive chromatin (4). Therefore, H3K36me2 maintains gene expression by protecting the genomic regions from the spreading of the repressive chromatin domain. Moreover, H3K36me2 serves as a binding site for DNMT3A, a *de novo* DNA methyltransferase, thereby controlling the DNA methylation pattern mainly in the intergenic regions (5).

Several lines of evidence have demonstrated the critical importance of strict regulation of cellular H3K36me2 levels. First, haploinsufficiency of NSD2 or NSD1 is known to be involved in Wolf–Hirschhorn or Sotos syndromes, respectively (6, 7). Second, aberrant upregulation of cellular H3K36me2 levels, induced by the overexpression or point mutation of NSD2, have been found in various cancers. Approximately 15–20% of patients with multiple myeloma carry a t(4;14) translocation, which causes the immunoglobulin heavy chain locus *(IGH)-NSD2* hybrid gene to induce overexpression of NSD2, along with a global increase and redistribution of H3K36me2 in the affected cells (8). Increased H3K36me2 reprograms the cells by reversing the repressive function of PRC2 in the myeloma (9). A recurrent point mutation, NSD2 p.E1099K in the catalytic SET domain, has been found in patients with acute lymphoblastic leukemia (10) and other types of cancers (11), and is known to aberrantly activate the H3K36 methyltransferase activity (10, 11). Another recurrent mutation, NSD2 p.T1150A in the SET domain, has been found in mantle cell lymphoma along with the p.E1099K mutation (12); however, to our knowledge, its effect on the catalytic activity has not yet been reported.

Biochemical studies have revealed that nucleosome structures regulate the methylation activity of NSD proteins. NSD2 exhibits weak and non-specific lysine methylation activity on histone octamer substrates, whereas it strongly and specifically methylates H3K36 when nucleosomal substrates are used (13). The linker histone H1, on the other hand, inhibits the activity of NSD2 on nucleosomal substrates (14). Crystal structures of the SET domains of NSD1 (15), NSD2 (16), and NSD3 (PDB 5UPD) have shown the substrate-binding cleft to be occupied by the autoinhibitory loop (residues 1180-1188 of NSD2), preventing H3K36 binding. Therefore, how NSD2 engages with nucleosomal H3K36 and how the oncogenic mutations alter its catalytic activity have been elusive.

## Results and discussion

### Overall structure of the NSD2-nucleosome complex

To understand how NSD2 engages with nucleosomal H3K36 and how the oncogenic mutations affect its catalytic activity, we solved a 2.8-Å resolution cryo-electron microscopy (cryo-EM) structure of the complex formed by the catalytic fragment of human NSD2 p.E1099K (residues 973–1226, containing the AWS domain, SET domain, and C-terminal basic extension) and nucleosome, in the presence of sinefungin, an analog of *S*-adenosyl methionine (SAM) (Supplementary Fig. S1–S3, Supplementary Table 1). To facilitate complex formation, we created an NSD2-H4 fusion protein connected by a 32-residue linker and co-expressed it with H3. We then assembled a nucleosome using the co-expressed proteins, along with H2A, H2B, and a 185-bp DNA possessing the 601 nucleosome positioning sequence at the center and a 20-bp linker DNA at both ends. Additionally, we introduced another substitution H3K36M, which is found in most patients with chondroblastoma (17) and is known to inhibit several H3K36 methyltransferases (18).

Figure 1 shows the overall structure of the NSD2-nucleosome complex. Although one nucleosome should contain two copies of NSD2 (due to the NSD2-H4 fusion used), we observed only one NSD2 molecule engaging with one of the two H3K36M residues; the other one possibly exhibits random orientation with respect to the nucleosome position. Residues 986– 1203 of NSD2 and 31–134 of the engaged H3 are visible in cryo-EM density and have been included in the model coordinates. The basic C-terminal extension (residues 1209–1226) is required for efficient nucleosome H3K36 methylation by NSD proteins (19). Although no clear density was observed, the extension is expected to be located near the nucleosomal DNA and form ionic interactions with the DNA phosphates. Upon nucleosome binding, NSD2 was seen to change its conformation markedly at the autoinhibitory loop to allow accommodation of the H3 tail (see “**Conformational change of the autoinhibitory loop**”).

**Figure 1.**
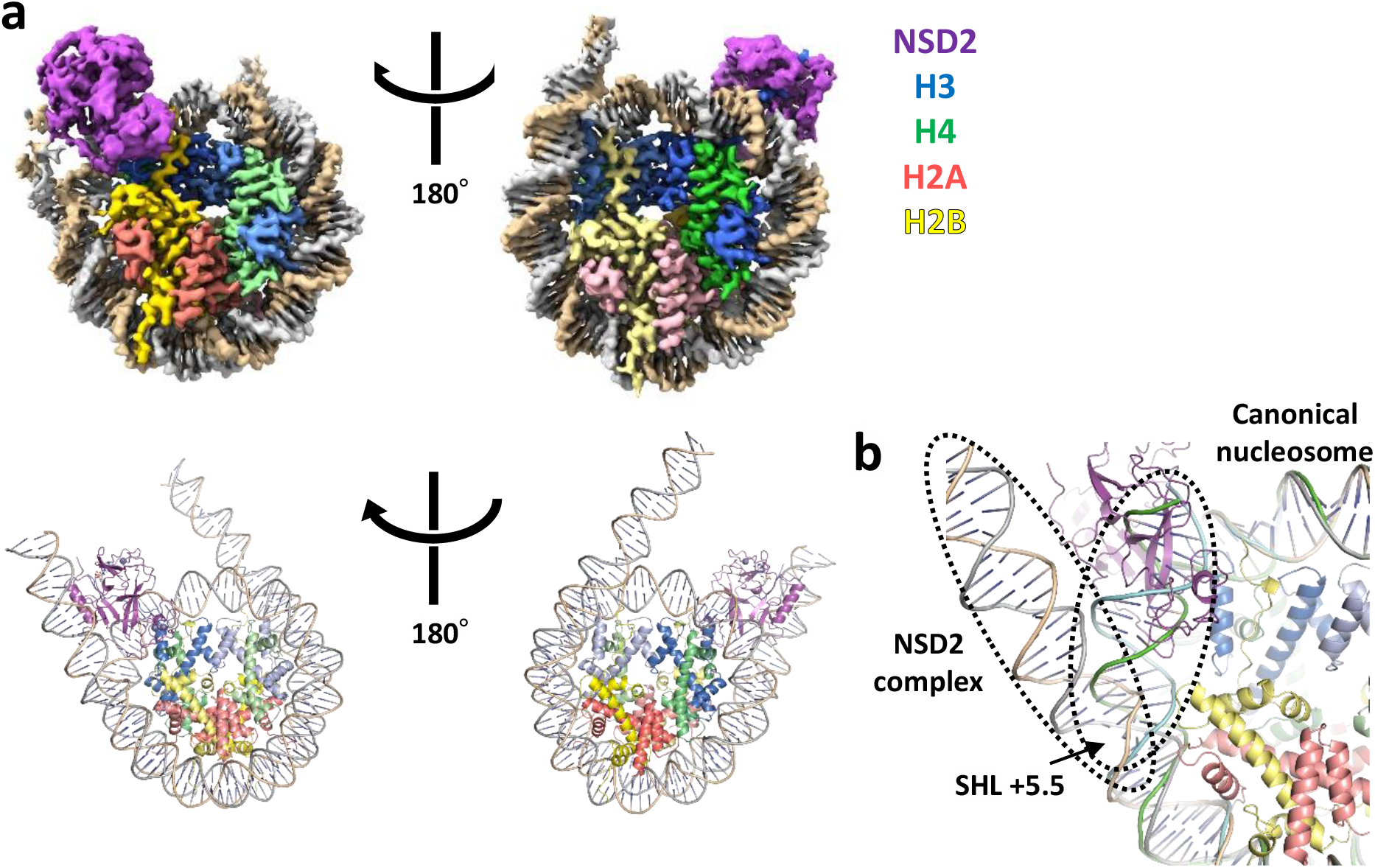
Overall structure. Unless stated otherwise, proteins are colored as follows: NSD2, magenta; H3, blue; H4, green; H2A, pink; and H2B, yellow. **a**, Density map (top) and ribbon model (bottom), two views related by 180° rotation. **b**, Superposition of the current NSD2-nucleosome structure with that of a canonical nucleosome (PDB 1KX5). Duplex DNA in the NSD2 complex stretches straight up to approximately SHL +5.5, resulting in its partial unwrapping.

A striking feature of the complex is the partial unwrapping of nucleosomal DNA near the entry site (Fig. 1a, b). In the canonical nucleosome, H3K36 is located between the two gyres of DNA in a relatively crowded environment. Thus, NSD2 needs to remove one gyre to gain access to H3K36 and accommodate its side chain into the catalytic pocket. The nucleosomal DNA, which usually exhibits a curved structure wrapping around the histone octamer, stretches around the superhelix location (SHL) +5.5, and extrudes away from the histone octamer. This extrusion allows NSD2 to establish interactions with the first α-helix of H3, which interacts with DNA in the canonical nucleosome. Moreover, NSD2 interacts with the N-terminal tail of H3, C-terminal portion of H2A, and DNA at two locations, SHL-1 and the external linker DNA position (Fig. 2a and Supplementary Fig. S4). The interaction modes are similar to those observed in the complex between yeast Set2, an H3K36 tri-methyltransferase, and nucleosome (20). However, the linker DNA in the two complexes exhibit different conformations, and three lysine residues of NSD2 (K992, K995, and K998), conserved across NSD proteins but not in Set2, face the linker DNA (Supplementary Fig. S3).

### Relationship between nucleosome structure and H3K36 methylation

The linker histone H1 has been shown to inhibit the methyltransferase activity of NSD2 toward chromatin (14). H1 binds to the nucleosome dyad and contacts with both linker DNAs, stabilizing a compact conformation (21, 22) (Supplementary Fig. S4). H1 may thus inhibit the activity of NSD2 by hindering the unwrapping of nucleosomal DNA and, hence, its access to H3K36.

Several chromatin factors, such as ATP-dependent chromatin remodelers (23), so-called “pioneer transcription factors,” which can recognize chromatinized DNA elements (24), and RNA polymerase II (25), are known to affect the conformation of nucleosomal DNA. Interestingly, their binding to nucleosomes resulted in the partial unwrapping of nucleosomal DNA, similar to that in the NSD2 complex (Supplementary Fig. S5). It may be speculated that the chromatin factors and H3K36 methyltransferases work cooperatively, so that the H3K36 exposure caused by other factors (such as chromatin transcription by RNA polymerase II) facilitates efficient methylation. In that case, H3K36 is well positioned to act as a memory mark for changes in the nucleosomal structure, as conformational changes of nucleosomal DNA affect its accessibility.

### Interactions of NSD2 with histones and DNA

Figure 2 shows the interactions between NSD2 and histones in detail. Two arginine residues on the first α-helix of H3 (H3R49 and H3R52) make extensive contacts with NSD2 (Fig. 2a). In the canonical nucleosome, these arginine residues are located close to the phosphate groups of DNA. The interactions between NSD2 and the two arginine residues may compensate for the loss of their interactions with DNA. In addition, H3Y41 stacks against K1152, which further forms a salt bridge with the phosphate group of nucleosomal DNA at SHL-1.

**Figure 2.**
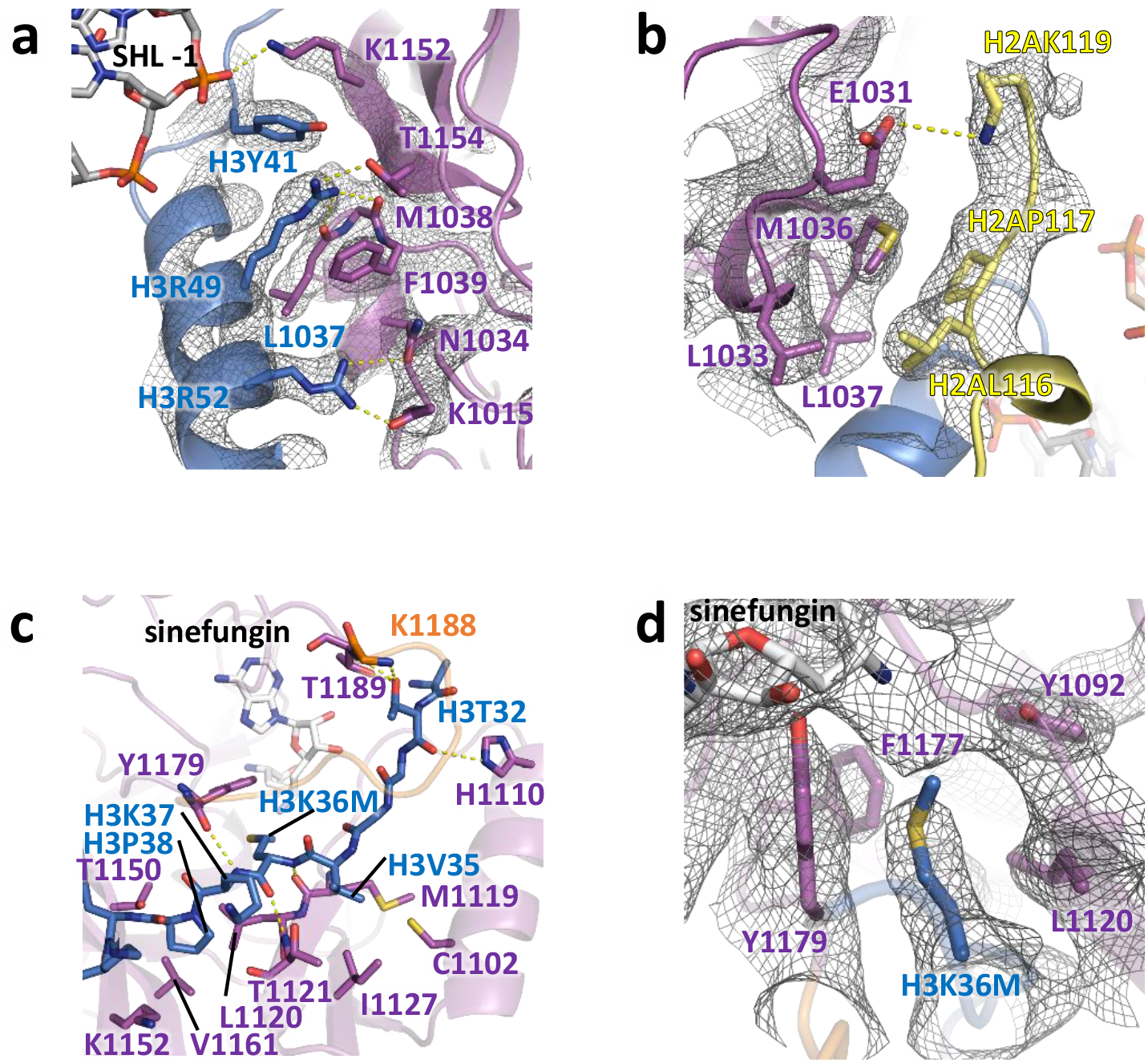
Interactions of NSD2 with histones H3 and H2A, and DNA. The cryo-EM density is shown as gray mesh in **a** (contoured at 7σ), **b** (at 3σ), and **d** (at 5σ). **a**, Interactions with the first α-helix of H3 and DNA. **b**, Interactions with H2A. **c**, Interactions with the H3 tail region. **d**, The H3K36- binding cavity.

The interactions between NSD2 and H2A are mainly hydrophobic, and there is a salt bridge between E1031 and H2AK119 (Fig. 2b). H2AK119 is mono-ubiquitinated by polycomb repressive complex 1 (PRC1), which cooperates with PRC2 to establish a transcriptionally repressive chromatin environment (26). Given the known antagonism between H3K36 methylation and polycomb group proteins, it would be interesting to examine whether H2AK119 mono-ubiquitination suppresses the H3K36 dimethylation activity of NSD2.

H3K36M and adjacent H3 residues are bound to the substrate-binding cleft on the surface of the SET domain of NSD2 (Fig. 2c). H3K36M and H3K37 form an intermolecular three-stranded β-sheet structure with M1119-T1121 on one side and Y1179 on the other. The H3T32 carbonyl and hydroxyl groups also make hydrogen bonds with NSD2. Two H3 residues, H3V35 and H3P38, are bound to small hydrophobic patches on the surface of NSD2. The H3V35 side chain interacts with C1102, M1119, T1121, and I1127, while the H3P38 side chain interacts with L1120, T1150, and V1161. The H3K36M side chain is accommodated in the catalytic cavity and surrounded by side chains of Y1092, L1120, F1177, and Y1179 (Fig. 2d). Structural comparison with the human SETD2-H3K36M peptide complex (27) revealed the aforementioned three aromatic residues to be conserved, whereas L1120 is replaced with methionine in SETD2 (Supplementary Fig. S3, S6). The different residues at this position may be related to the difference in the methylation state specificity between the NSD family (dimethyltransferases) and SETD2 (a trimethyltransferase).

### Conformational change of the autoinhibitory loop

Figure 3 shows the structural comparison between the H3-free NSD2 (16) and the current NSD2 engaging nuclesomal H3K36. In the H3-free NSD2, the autoinhibitory loop (N1180-K1188) adopts a closed conformation, occupying the substrate-binding cleft. Upon nucleosome binding, the loop moves to adopt an open conformation, making room for the binding of H3K36M and nearby residues.

**Figure 3.**
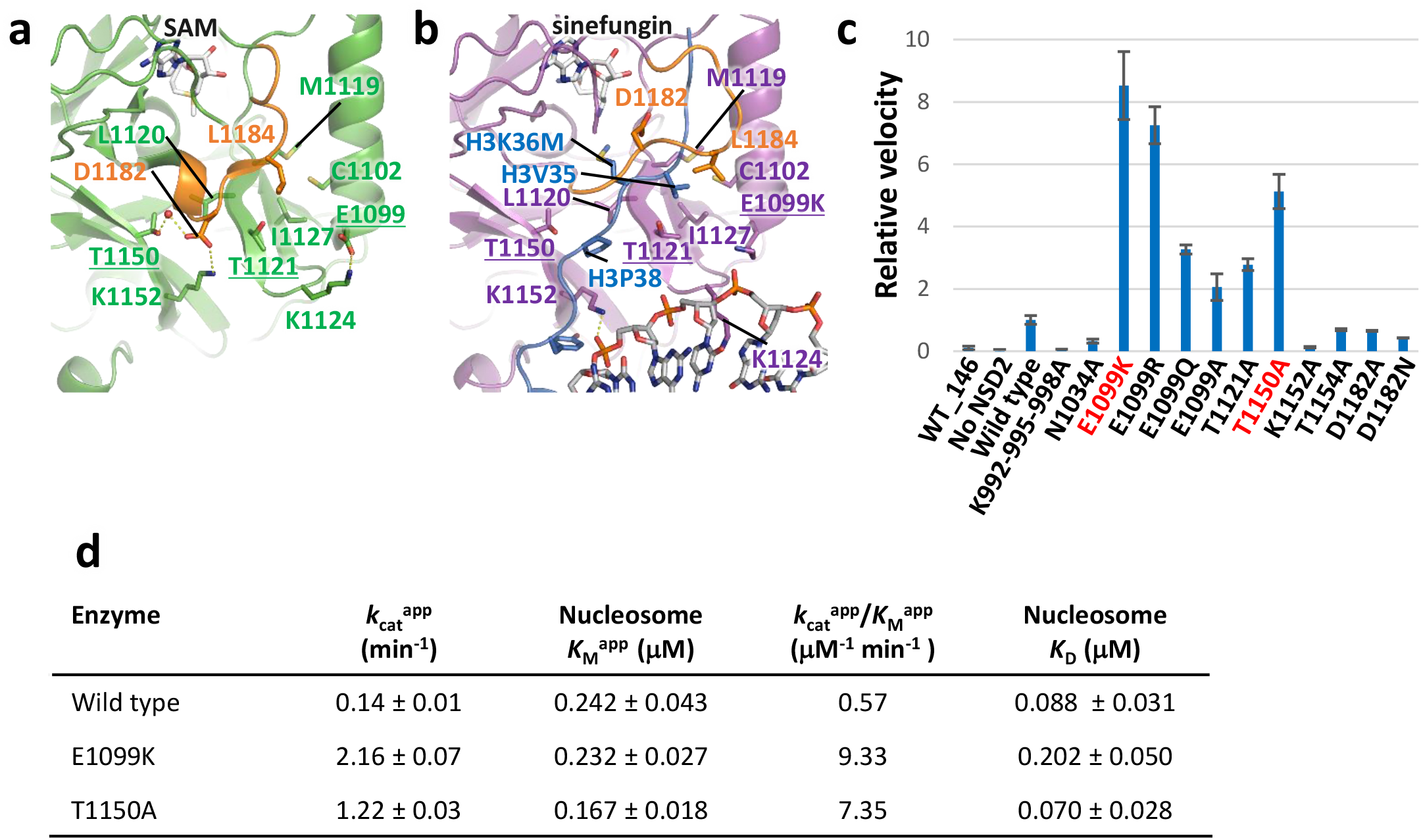
Autoinhibitory loop and oncogenic mutations. **a** and **b**, Structures of the H3-free NSD2 (**a**) and NSD2-nucleosome complex (**b**). Residues whose substitutions resulted in increased H3K36 methyltransferase activity are shown with underlined labels. The autoinhibitory loop is colored orange. A water molecule mediating the interaction with T1150 and D1182 in the H3-free form is shown as a red sphere. **c**, Summary of the enzymatic analysis, showing the relative reaction velocity. **d**, Kinetic values and dissociation constants between the NSD2 proteins and the nucleosome with 185-bp DNA.

Detailed structural comparison revealed two elements that could be important for such conformational transition. First, in the H3-free form, D1182 of the autoinhibitory loop forms a network of water-mediated hydrogen bonds and a salt bridge with side chains of T1150 and K1152, apparently stabilizing the loop conformation (Fig. 3a). Similar interactions are also found in the crystal structure of NSD1 (28), implying the functional importance of this conformation shared across the NSD family proteins. The T1150A substitution, recurrently found in mantle cell lymphoma (12), would aberrantly increase the catalytic activity of NSD2 by disrupting the interaction with D1182, thereby affecting the conformation of the autoinhibitory loop. In the nucleosome complex, K1152 of NSD2 no longer forms a salt bridge with D1182, and rather interacts with H3Y41 and a DNA phosphate (Fig. 2a, 3b). Thus, K1152 may couple the movement of the autoinhibitory loop with nucleosome binding by switching its binding partner in these forms. This mechanism is also consistent with the previous report that the addition of 41-bp DNA was shown to stimulate the catalytic activity of NSD1 and NSD2 on histone octamers (13).

Second, in the H3-free form, the side chain of L1184 in the autoinhibitory loop binds to the hydrophobic patch formed by C1102, M1119, T1121, and I1127 (Fig. 3a). In the nucleosome complex, the same patch binds to the H3V35 side chain (Fig. 3b). The oncogenic E1099K mutation site is located close to the patch. In the H3-free form, the E1099 side chain forms a salt bridge with K1124, which is located on a loop connecting the two β-strands harboring M1119, T1121, and I1127 (Fig. 3a). Thus, E1099K substitution would disrupt the salt bridge with K1124, possibly affecting the conformational flexibility of the hydrophobic patch and eventually that of the autoinhibitory loop.

### E1099K and T1150A increase the apparent *kcat* value of NSD2

To gain insight into the mechanisms of substrate recognition and their dysregulation by oncogenic mutations, we conducted biochemical analyses (Fig. 3c). First, we measured the nucleosomal H3K36 methyltransferase activity of NSD2 and its mutants using a commercial kit that couples the production of *S*-adenosyl homocysteine (SAH), a reaction byproduct, to chemiluminescence (29). NSD2 efficiently methylated the nucleosomes with 185-bp DNA, though not those with 146-bp DNA, suggesting the importance of its interaction with a linker DNA. Consistently, when the three lysine residues (K992, K995, and K998) that face the linker DNA (Supplemental Fig. S4) were all mutated to alanine, the mutant lost its catalytic activity. The N1034A, K1152A, and T1154A substitutions reduced the activity, showing the important roles played by the interactions of the residues with the nucleosome (Fig. 2a, Supplementary Fig. S3).

We next confirmed that the E1099K mutant has a stronger catalytic activity than the wild-type (Fig. 3c). Additionally, substituting E1099 with arginine, glutamine, and alanine resulted in stronger activities (Fig. 3c), suggesting the essential regulatory role of glutamate at this position. Interestingly, another oncogenic mutant, T1150A, also exhibited stronger activity (Fig. 3c). Moreover, we found that the T1121A substitution, which alters one of the H3V35-binding patch residues (Fig. 3b), stimulates the catalytic activity (Fig. 3c). This structure-guided identification of a novel activating mutation suggests an important role of the H3V35-binding patch for the autoinhibition.

Next, we conducted a kinetic analysis to gain further insight into the dysregulated oncogenic mutations (Fig. 3d, Supplementary Fig. S7). The results showed the increase in catalytic efficiency (*k*_cat_^app^W_M_^app^) to be mainly governed by an increase in the apparent catalytic turnover (*k*_cat_^app^) value for both E1099K and T1150A mutants (approximately 16- and 9-fold, respectively). The differences in the *k*_M_^app^ values toward nucleosomes were small across the wild-type, E1099K, and T1150A proteins. We also measured the affinity between NSD2 and nucleosomes using microscale thermophoresis experiments (30), which showed that neither E1099K nor T1150A increased the nucleosome affinity (Fig. 3d, Supplementary Fig. S8). The E1099K mutant exhibited even weaker affinity, although the positively charged K1099 side chain is near the nucleosomal DNA. Taken together, the oncogenic E1099K and T1150A mutations both increased the catalytic turnover, though not the nucleosome affinity, of NSD2.

### Impact of E1099K and T1150A on the dynamics of the autoinhibitory loop

To investigate the impact of E1099K and T1150A mutations on the dynamics of the autoinhibitory loop, we performed MD simulations of the SET domain of the NSD2-SAM complex in the absence of H3, starting from the current nucleosome-engaging structure. We performed runs on the wild-type and three NSD2 mutants (E1099K, T1150A, and the E1099K-T1150A double mutant, Supplementary Table 2) and analyzed the trajectories obtained from three independent 500-ns runs for each system. In all the systems, the conformation of the autoinhibitory loop was flexible during the simulation, suggesting that the H3-engaging conformation is not stable in the absence of H3 (Supplementary Movie 1).

We first calculated the dynamic cross-correlation matrix of the SET-domain residue fluctuations to capture the correlated motions among the residues. In the wild-type, we observed anti-correlated motions between residues 1095–1130 (region R1), 1140–1150 (region R2), as well as R1 and residues 1177–1203 (region R3) (Fig. 4a, b). R1 contained E1099 and residues forming the H3V35- and H3P38-binding hydrophobic patches, whereas R2 and R3 contained T1150 and the autoinhibitory loop, respectively. In all three mutants, the anti-correlated motions between R1– R2 and R1–R3 disappeared (Fig. 4a), indicating that the movement of R2 and R3 were independent of R1.

**Figure 4.**
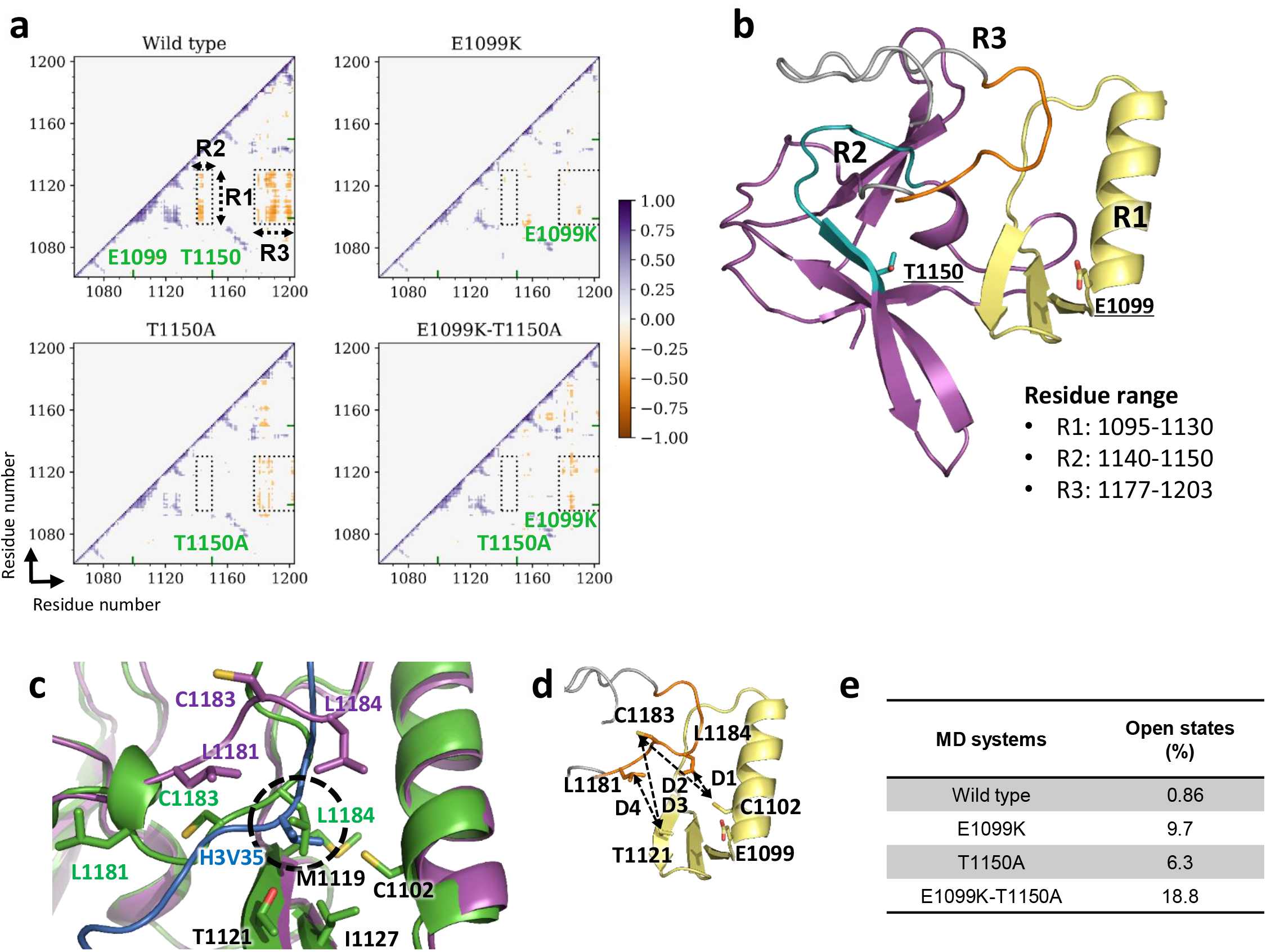
Opening and closing of the autoinhibitory loop. **a**, 2D plots of the residue-wise cross-correlation matrices of NSD2 wild-type, E1099K, T1150A, and double mutant E1099K-T1150A. All the correlation coefficients between −0.3 and 0.3 were taken as zero to visualize the significantly correlated residues. The regions labeled by R1 (residues 1095– 1130), R2 (residues 1140-1150), and R3 (PSL) exhibit a remarkable difference between the wild-type and mutants. The mutation sites are denoted by green ticks on the axis. **b**, The regions R1 (pale yellow), R2 (teal), and R3 (gray and orange), as well as the mutation sites (stick view), are shown and labeled on the NSD2 SET domain structure. **c**, Superimposition of the H3-free NSD2 (green) and nucleosome-bound NSD2 (magenta). H3 is shown in marine blue. The overlapping hydrophobic cavity occupied by the L1184 residue in substrate-free NSD2 and H3V35 is encircled in black. **d**, Definition of four distance-based locks (D1: L1184-C1102, D2: C1183- C1102, D3: C1183-T1121, and D4: L1181-T1121), as represented by dashed lines on SET domain structure. The autoinhibitory loop is shown in orange and residues are labeled. **e**, Table showing the percentage of open autoinhibitory loop conformations observed under different conditions.

The independent movement of R3 suggested that the autoinhibitory loop might tend to adopt conformations that allow histone H3 binding in the mutants more often. To test this hypothesis, we considered four distance-based locks (D1, D2, D3, and D4) that represent the hydrophobic interactions between residues on the autoinhibitory loop (L1181, C1183, and L1184) and those in the H3V35-binding patch (C1102 and T1121). We defined the autoinhibitory loop as “open” when all the four locks are released, and “closed” otherwise. The probabilities of open states of the loop and individual locks are shown in Fig. 4e and Supplementary Table 3. In the wild-type, the autoinhibitory loop was open only for 0.86% of the simulation time, indicating that it almost always occupies the substrate-binding cleft. In the E1099K, T1150A, and E1099K-T1150A mutants, open states appeared far more frequently (9.7%, 6.3%, and 18.8%, respectively). A detailed examination of the MD trajectories revealed a novel mode of interaction between the autoinhibitory loop and the hydrophobic patches (Supplementary Fig. S9). In the wild-type, the L1181 side chain often forms hydrophobic interactions with T1121 and/or T1150, the two threonine residues whose substitutions lead to aberrant enzymatic activation. These interactions occurred less often in the three mutants, as indicated by the larger chances of the released state of D4 lock. Collectively, our MD analysis suggests that the E1099K and T1150A mutations affect the dynamics of autoinhibitory loop, causing NSD2 to adopt open states more often to accommodate the H3 tail in the substrate-binding cleft.

### A model for autoinhibition and its dysregulation by E1099K and T1150A

Based on our analyses, we propose a mechanistic model in which the autoinhibitory loop conformation regulates the catalytic activity of NSD2, and the oncogenic E1099K and T1150A mutations aberrantly disturb the autoinhibition (Fig. 5). In the H3-free form, the autoinhibitory loop dynamically moves and exhibits multiple conformations (two possible major conformations are indicated in Fig. 5) while remaining bound to the H3V35- and H3P38-binding patches by hydrophobic interactions via L1181, C1183, and L1184, and by hydrophilic interactions via D1182. Consequently, the substrate-binding cleft is almost always occupied by the autoinhibitory loop, leaving no room for H3 binding. Upon nucleosome binding, the conformation of autoinhibitory loop changes, allowing H3V35 and H3P38 to be accommodated in the hydrophobic patches and enabling precise positioning of the H3K36 side chain for methyl group transfer. This transition is partially triggered by the binding of H3Y41 and nucleosomal DNA to K1152, which is incompatible with its interaction with D1182. In the E1099K mutant, the salt bridge formed between E1099 and K1124 is lost, affecting the local conformation or dynamics of the H3V35-binding patch. This, in turn, could affect the binding of C1183 and L1184 in the autoinhibitory loop to the H3V35-binding patch, resulting in its higher tendency to adopt open states and an increase of catalytic turnover. Similarly, T1150A increases open states by disrupting the hydrophilic interaction with D1182 and/or the hydrophobic interactions with L1181 in the autoinhibitory loop. It will be interesting to examine whether a similar mechanism regulates the activity of other H3K36 methyltransferases such as NSD1, NSD3, and SETD2.

**Figure 5.**
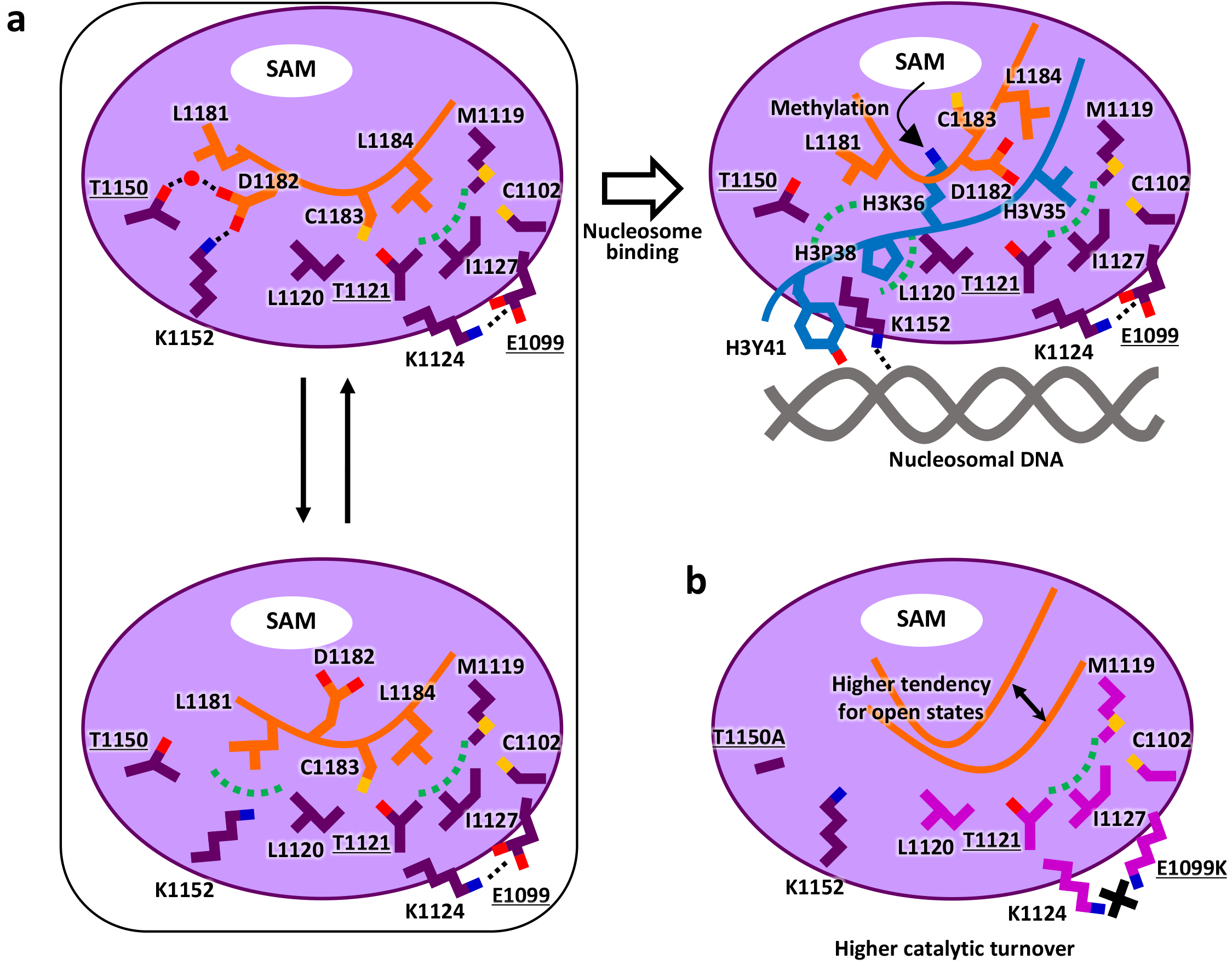
Mechanistic model of how the conformational dynamics of the autoinhibitory loop regulates the catalytic activity of NSD2. **a**, Normal regulation. In the H3-free state, though the autoinhibitory loop dynamically moves (two possible conformations are indicated on the left panels), it remains bound to the substrate-binding cleft by interactions around two hydrophobic patches. Upon nucleosome binding (the upper right panel), the autoinhibitory loop moves to make room for H3 binding. The structural transition is partly triggered by the interaction of K1152 with H3Y41 and DNA. **b**, Dysregulation by oncogenic E1099K and T1150A mutations. E1099K disrupts the salt bridge between E1099 and E1124, affecting the local conformation or dynamics of the H3V35-binding patch (indicated by a lighter magenta color of the affected residues) and its interactions with the autoinhibitory loop. T1150A disrupts the hydrophobic or hydrophilic interactions between the H3P38-binding patch and the autoinhibitory loop. Both mutations, thus, result in a higher tendency to adopt open states and increase the catalytic turnover.

NSD2 is implicated in various types of cancers, including multiple myeloma and acute lymphoblastic leukemia (31). Our study should guide the development of specific inhibitors for the treatment of these cancers. Moreover, multiple histone lysine methyltransferases are known to be under complex regulation, involving, allosteric regulators, and post-translational modifications (32–35). Our detailed analysis and the proposed mechanistic model may help understand the dynamic regulatory mechanisms potentially employed by other lysine methyltransferases that use the SET domain for catalysis.

## Methods

### DNA preparation

We prepared 146-bp and 185-bp DNA fragments using different strategies. The 146-bp DNA fragment was excised from a plasmid DNA containing 14 copies of the Widom 601 DNA sequence by EcoRV. The excised DNA was purified by HiTrap Q column chromatography (GE Healthcare), flash-frozen with liquid nitrogen (LN_2_), and stored at −80 °C. The 185-bp DNA fragment was generated using nested-PCR amplification. Initially, a 282-bp DNA fragment containing the Widom 601 DNA sequence was amplified using pGEM-3Z-601 (36) as template and 282F (5′-CGGGATCCTAATGACCAAGGAAAGC-3’) and 282R (5′-GGGAGCTCGGAACACTATCCGAC-3’) as primers, and subsequently purified by acrylamide gel electrophoresis. The 185-bp DNA fragment was amplified with the 282-bp DNA fragment as template and 185F (5′-GACCCTATACGCGGCCGCCCTGG-3’) and 185R (5′-GTCGCTGTTCAATACATGCACAGGATG-3’) as primers. The amplified 185-bp DNA fragment was purified by HiTrap Q column chromatography flash-frozen with LN2, and stored at −80 °C.

### Protein preparation

Human histone H2A was bacterially expressed as a His6- and SUMOstar-tagged protein, and isolated from inclusion bodies under denaturing conditions (20 mM Tris-HCl pH 7.5, 7 M guanidine hydrochloride, 500 mM NaCl, and 1 mM DTT). The supernatant was purified using Ni-resin and dialyzed against a buffer containing 20 mM Tris-HCl pH 7.5, 150 mM NaCl, and 0.5 mM DTT. The His6-SUMOstar-tag was removed by SUMOstar protease digestion. The resulting histone H2A was further purified by HiTrap SP column chromatography (GE Healthcare), flash-frozen with LN2, and stored at −80 °C.

Human histone H2B was bacterially expressed as a His6-tagged protein, and isolated from inclusion bodies under a denaturing condition (20 mM Tris-HCl pH 7.5, 7 M guanidine hydrochloride, 500 mM NaCl, and 1 mM DTT). The supernatant was purified using Ni-resin and dialyzed against a buffer containing 20 mM Tris-HCl pH 7.5, 100 mM NaCl, and 1 mM DTT. The His6-tag was removed by thrombin digestion. The resulting histone H2B was further purified by HiTrap SP column chromatography, flash-frozen with LN2, and stored at −80 °C.

His6 and SUMOstar-tagged human histone H4 was bacterially co-expressed with human histone H3 and purified under non-denaturing conditions. Cells co-expressing H3 and H4 were suspended in a buffer containing 40 mM K2HP04, 10 mM KH2P04, 20 mM imidazole, 3 M NaCl, 1 mM DTT, 0.1 mM PMSF, and 0.5% TritonX-100, and sonicated. The supernatant was purified using HisTrap column chromatography (GE Healthcare). The His6-SUMOstar-tag was removed by SUMOstar protease digestion. The resulting histone H3/H4 complex was further purified by HiTrap SP column chromatography, flash-frozen with LN2, and stored at −80 °C.

His6 and SUMOstar-tagged human NSD2-E1099K were fused to H4 and bacterially co-expressed with human histone H3 possessing K36M substitution. The complex between NSD2-E1099K-H4 and H3 was purified in a way similar to the H4-H3 complex; however, the His6-SUMOstar tag was not removed.

Wild-type and mutant NSD2 proteins (973-1226) containing Twin-Strep-tag and His6-tag at the N-terminus were bacterially expressed, purified using the StrepTrap column (GE Healthcare), flash-frozen with LN2, and stored at −80 °C.

### Histone octamer preparation

We incubated histones H2A, H2B, and H3/H4 at a molar ratio of 1.2:1.2:1 in the unfolding buffer (6 M guanidine hydrochloride, 4 mM HEPES-Na pH 7.5, 200 mM NaCl, and 5 mM DTT) at 4 °C for 1 h, followed by dialysis against refolding buffer (10 mM Tris-HCl pH 7.5, 2 M NaCl, 1 mM EDTA, and 5 mM 2-mercaptoethanol) to reconstitute the histone octamer. The latter was eventually purified from the unincorporated components using a Superdex 200 pg 26/60 column (GE Healthcare) equilibrated with buffer containing 10 mM Tris-HCl pH 7.5, 2 M NaCl, 1 mM EDTA, and 5 mM 2- mercaptoethanol, flash-frozen with LN2, and stored at −80 °C.

### Nucleosome reconstitution

To reconstitute nucleosomes used for enzymatic and interaction analyses, we mixed 185-bp (final 6.1 μM) or 146-bp DNA (final 5.6 μM) with a histone octamer at a molar ratio of approximately 1:1.1, and dialyzed the same against 125 mL of 10 mM HEPES-Na pH 7.5, 2 M KCl, and 1 mM DTT for 1 h. Thereafter, 875 mL of 10 mM HEPES-Na pH 7.5 and 1 mM DTT was gradually added to facilitate nucleosome reconstitution. The nucleosome samples were further dialyzed against 10 mM HEPES-Na pH 7.5, 50 mM KCl, and 1 mM DTT. The centrifuged supernatant was eventually concentrated and stored at 4 °C.

To prepare the NSD2-nucleosome complex for cryo-EM analysis, we mixed 185-bp DNA (final 2.4 μM) with H2A, H2B, and the complex between NSD2 E1099K-H4 and H3K36M at a molar ratio of approximately 1:2.6:2.6:2.6, and dialyzed the mixture for 1 h against 125 mL of 10 mM HEPES-Na pH 7.5, 2 M KCl, and 1 mM DTT solution. Thereafter, 875 mL of 10 mM HEPES-Na (pH 7.5) and 1 mM DTT was gradually added to facilitate the reconstitution of nucleosomes. The NSD2-nucleosome complex was further dialyzed against 10 mM HEPES-Na pH 7.5, 1 mM DTT, and 25 mM or 50 mM KCl. The centrifuged supernatant was concentrated, and the sinefungin solution (25 mM) added (final 1.3 mM) before cryo-EM data acquisition.

### Sample vitrification and cryo-EM data acquisition

The reconstituted NSD2-nucleosome complex was applied to a freshly glow-discharged Quantifoil holey carbon grid (R1.2/1.3, 300 mesh, Cu/Rh grid for the sample with 25 mM KCl, and Au grid for the sample with 50 mM KCl), blotted for 4 s at 4 °C in 100% humidity, and plunge-frozen in liquid ethane using a Vitrobot Mark IV (Thermo Fisher Scientific). Grid images were obtained using a 300 kV Titan Krios G3i microscope (Thermo Fisher Scientific) equipped with a K3 direct electron detector (Gatan) installed at the University of Tokyo, Japan. Data sets were acquired with the SerialEM software, with a defocus range of −0.8 to −1.6 μm. Data acquisition statistics are shown in Supplementary Table 1.

### Image processing and model building

Movie stacks were corrected for drift- and beam-induced motion using MotionCor2 (37), and the CTF parameters were estimated using GCTF (38). Particles were automatically picked using RELION-3.1 (39), and then used for two-dimensional (2D) classification, *ab-initio* reconstruction, and hetero refinement with cryoSPARC (40). The particles that converged to the NSD2-nucleosome complex class were exported to RELION-3.1, and used for the focused three-dimensional (3D) classification with a mask covering only NSD2. The particles that converged to the class with a well-resolved NSD2 density were used for final 3D refinement with RELION-3.1. The resolution was estimated based on the gold-standard Fourier shell correlation (FSC) curve at 0.143 criterion. The atomic models of H3-free NSD2 (PDB 5LSU) and nucleosome (PDB 1KX5) were fit into the density using UCSF Chimera (41), and then manually modified using Coot (42). Real-space refinement was performed using Phenix (43).

### Methyltransferase assay

For a simple comparison of the initial reaction rate, 128 nM wild-type or mutant NSD2, and 4 μM nucleosomes were used (Fig. 3c). For kinetic analysis, 256 nM wild-type NSD2, 16 nM E1099K mutant or 32 nM T1150A mutant, and a series of two-fold diluted nucleosome (from 31.25 nM to 4 μM) were used (Fig. 3d, Supplemental Fig. S7). NSD2 proteins, 30 μM SAM, and nucleosome were mixed in a reaction buffer (2.5 mM HEPES-Na pH 7.5, 10 mM Tris-HCl pH 9.5, 2.5 mM NaCl, 12.5 mM KCl, 1 mM TCEP, and 0.01% Tween 20), and incubated at 30 °C. Approximately 4 μL of each reaction mixture was taken at 2, 4, 6, and 8 min, and quenched by adding 1 μL of 0.5% TFA. Methyltransferase activity was evaluated using an MTase-Glo Methyltransferase Assay Kit (Promega). The luminescent signal corresponding to SAH production was measured using Centro LB 960 (Berthold Technologies) in a white half-area 96-well plate. The measured luminescent signal was converted to represent the amount of SAM utilized using a SAH standard curve. The initial rate of each reaction was determined using a linear regression fit of the data. Each reaction was run in triplicate. Kinetic parameters were derived by fitting the values to the Michaelis-Menten model using KaleidaGraph 4.5.3 software.

### Microscale thermophoresis

The microscale thermophoresis (MST) assay was performed using a Monolith NT.115 instrument (NanoTemper Technologies). For the MST assay, 50 nM His-tagged NSD2 proteins labeled with RED-Tris-NTA and series of two-fold diluted nucleosome or DNA (from 0.31 nM to 10 μM) were incubated in the binding buffer (20 mM HEPES-Na pH 7.5 150 mM KCl, 5 % glycerol, 0.05 % Tween 20, 1 mM DTT, and 1 mM sinefungin) for 30 min at room temperature, and centrifuged at 15000 rpm for 5 min. The samples were filled into premium capillaries to obtain measurements with an extinction power of 60% and medium MST power at 25 °C. Thermophoresis data were analyzed using MO.Affinity Analysis software ver. 2.3 (NanoTemper Technologies).

### Molecular dynamics simulations

The NSD2 E1099K structure (residues 986–1203 constituting two Zn atoms in the AWS domain, one Zn atom and SAM in the SET domain) was extracted from the cryo-EM-based nucleosome-bound NSD2 E1099K complex, in this study, by deleting the nucleosome. The N-terminal helix region (residues 973–985) from the crystal structure (16) (PDB ID: 5LSU) was added to the extracted NSD2 structure. In the combined structure, the L975 and L978 residues of N-terminal helix and L1071, Q1072, and R1073 residues of SET domain were mutated back to the original-sequence residues Q975, A978, D1071, G1072, and K1073, respectively, to obtain the initial structure of NSD2 E1099K with an open autoinhibitory loop conformation. Subsequently, the initial structures of NSD2 wild-type, and T1150A and E1099K-T1150A mutants were obtained by introducing appropriate mutations. All mutations were introduced using the rotamer library in UCSF Chimera (41, 44). Taken together, we performed MD simulations of NSD2 wild-type and three mutants, namely E1099K, T1150A, and E1099K-T1150A (Supplementary Table 2).

All the simulations were performed using the AMBER package with the ff14SB force field for protein and improved parameters for SAM (45–47). Parameters of the Zn ion coordinated to C1144, C1191, C1193, and C1198 were obtained from the zinc AMBER force field (ZAFF) (48). The parameters of the two Zn ions in the AWS domain, coordinated to seven cysteine residues, were generated using the MCPB.py program (49). Geometry optimization and RESP charge (Merz-Kollman scheme) calculations were performed at the B3LYP/6-31 * level of theory in Gaussian09 (50–52).

Each system was solvated and neutralized in a cubical box containing 0.150 M NaCl TIP3P water model *(53)* solution with a padding distance of 13.5 Å. Energy minimization involved both steepest-descent and conjugate-gradient algorithms. System equilibration was performed in three successive steps of 500 ps each, first, heating up to 300K in NVT, followed by equilibration under NPT at a temperature of 300 K and pressure of 1 bar. A positional restraint of 100 kcal mol^-1^ Å^-2^ was applied to the heavy atoms of the solute during the first two cycles of equilibration. The temperature at 300 K and pressure at 1 bar were maintained using the Langevin dynamics algorithm (collision frequency γ = 2.0 and coupling constant = 1.0 ps) and the Berendsen barostat (pressure relaxation time = 1.0 ps). Particle mesh Ewald (PME) was used to calculate the long-range electrostatic interactions (54), and the bonds associated with hydrogen atoms were constrained using the SHAKE algorithm (55). Finally, three independent 500-ns runs were performed for each system. All analyses of the trajectories were performed using the CPPTRAJ package (56).

### Figure preparation

The structural figures were created using PyMOL (Schrödinger, LLC) and UCSF ChimeraX (57). Sequence-alignment figure was created using ESPript (58). Both trajectory visualization and movie creation were accomplished using VMD (59, 60).

## Acknowledgements

Cryo-EM data were collected at the cryo-EM facility, University of Tokyo.

This research was supported by JSPS KAKENHI [grant numbers JP20K08717 (to K.S.), JP18H05534 (to H.K.), and JP20H05394 (to T.S.)]. The study was partially supported by the Platform Project for Supporting Drug Discovery and Life Science Research (Basis for Supporting Innovative Drug Discovery and Life Science Research (BINDS)) from AMED [grant numbers JP20am0101115 and JP20am0101106 (support numbers 1061 and 2450)].

## Author contributions

T.S. conceived the project. K.S., H.K., O.N., K.O., and T.S. designed the experiments. K.S., C.O., A.O., and T.S. cloned, expressed, and purified the proteins and prepared the nucleosomes. K.S., T.N., and T.S. collected and processed the cryo-EM data and built and refined the atomic model. K.S., K.H., C.O., A.M., and T.S. performed the biochemical experiments. A.K., S. S., and H.K. performed the MD simulations. K.S., A.K., H.K., K.O., and T. S. wrote the paper with input from all authors.

---

**Supplementary Figure S1.**
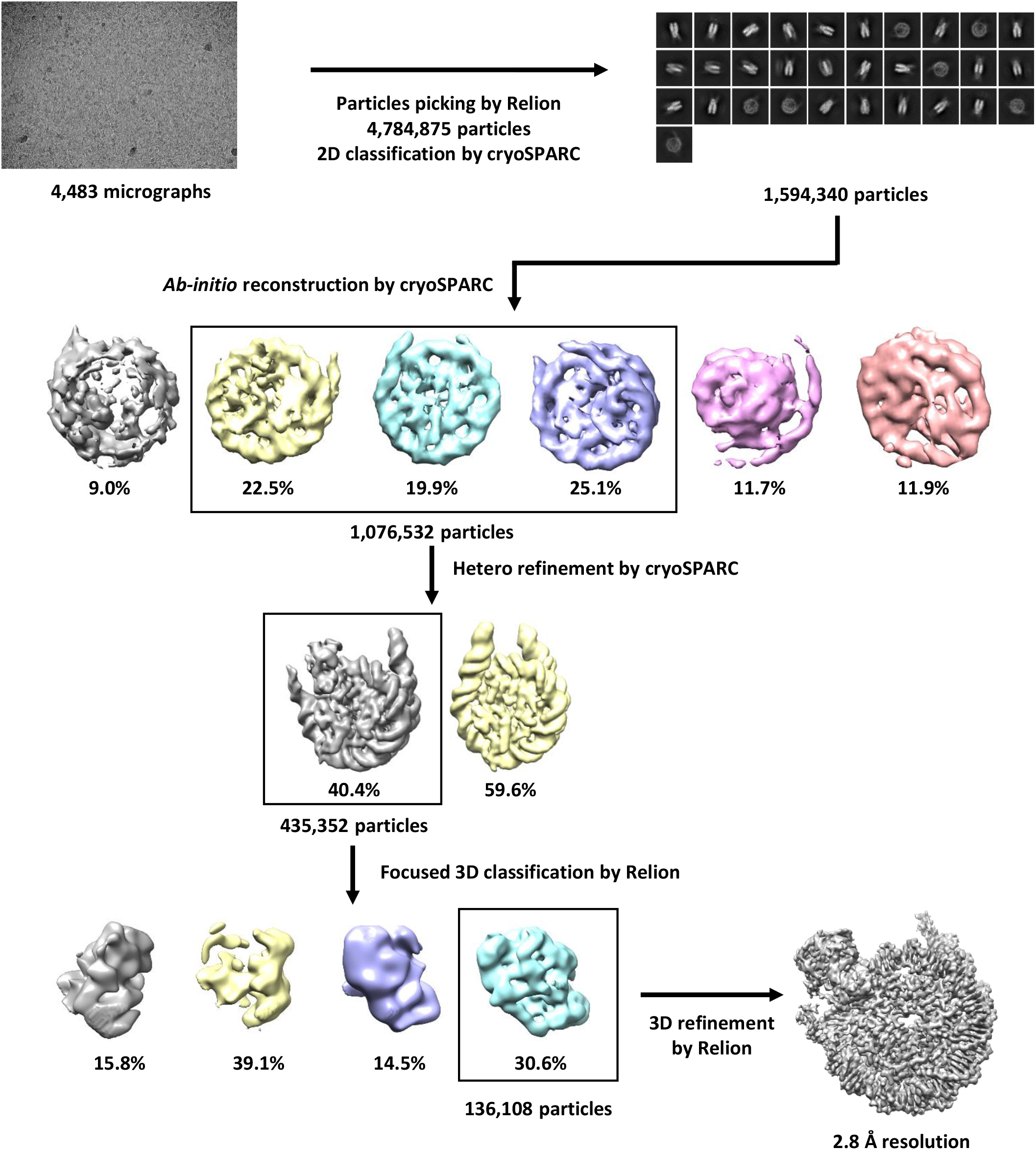
Cryo-EM 3D reconstruction and refinement.

**Supplementary Figure S2.**
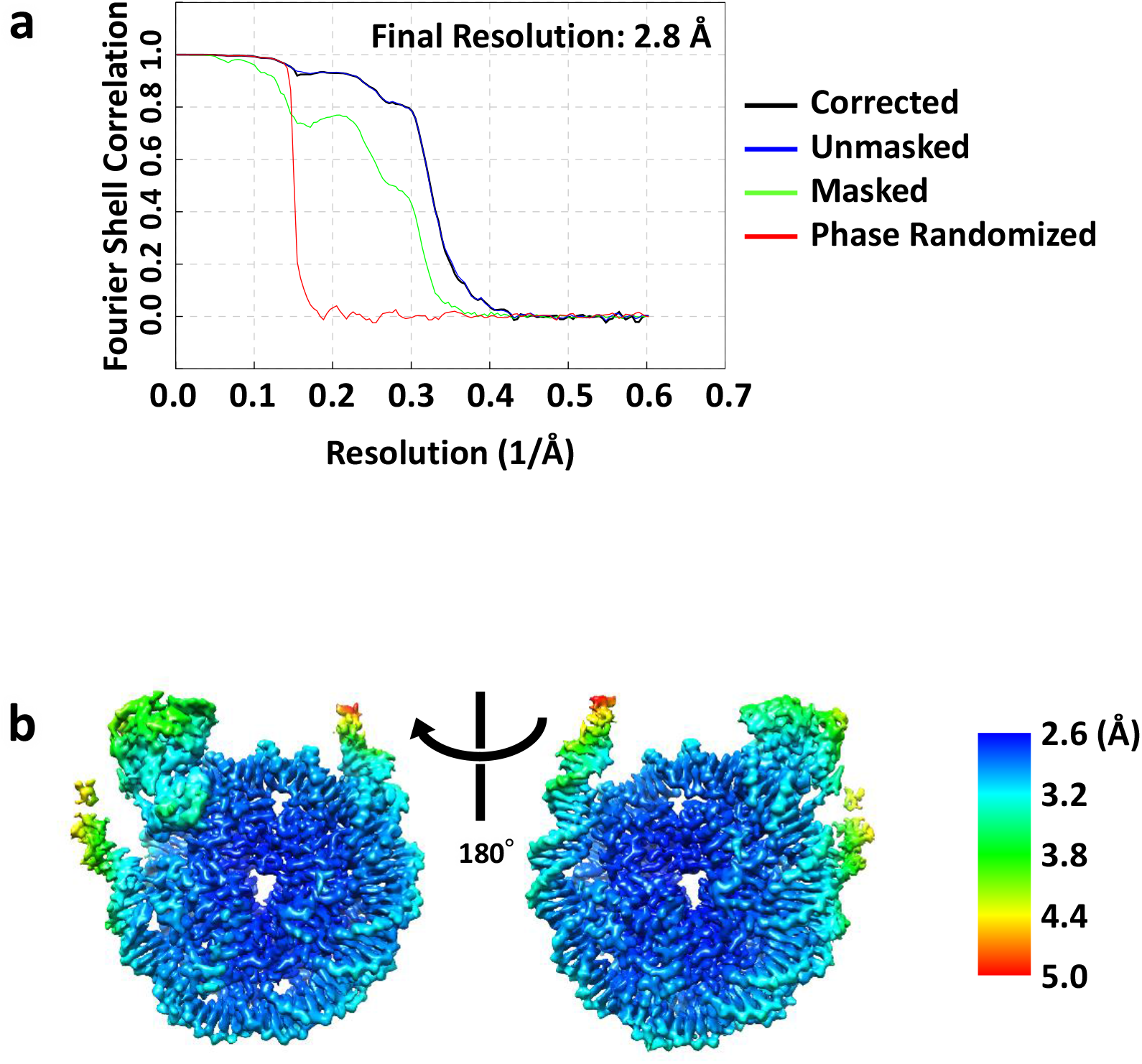
**Global** and local resolution of the 3D reconstituted map. **a**, Fourier shell correlation (FSC) curves of the 3D reconstruction. Resolution was estimated based on the FSC curve at 0.143 criterion. **b**, Local resolution map of the NSD2-nucleosome complex.

**Supplementary Figure S3.**
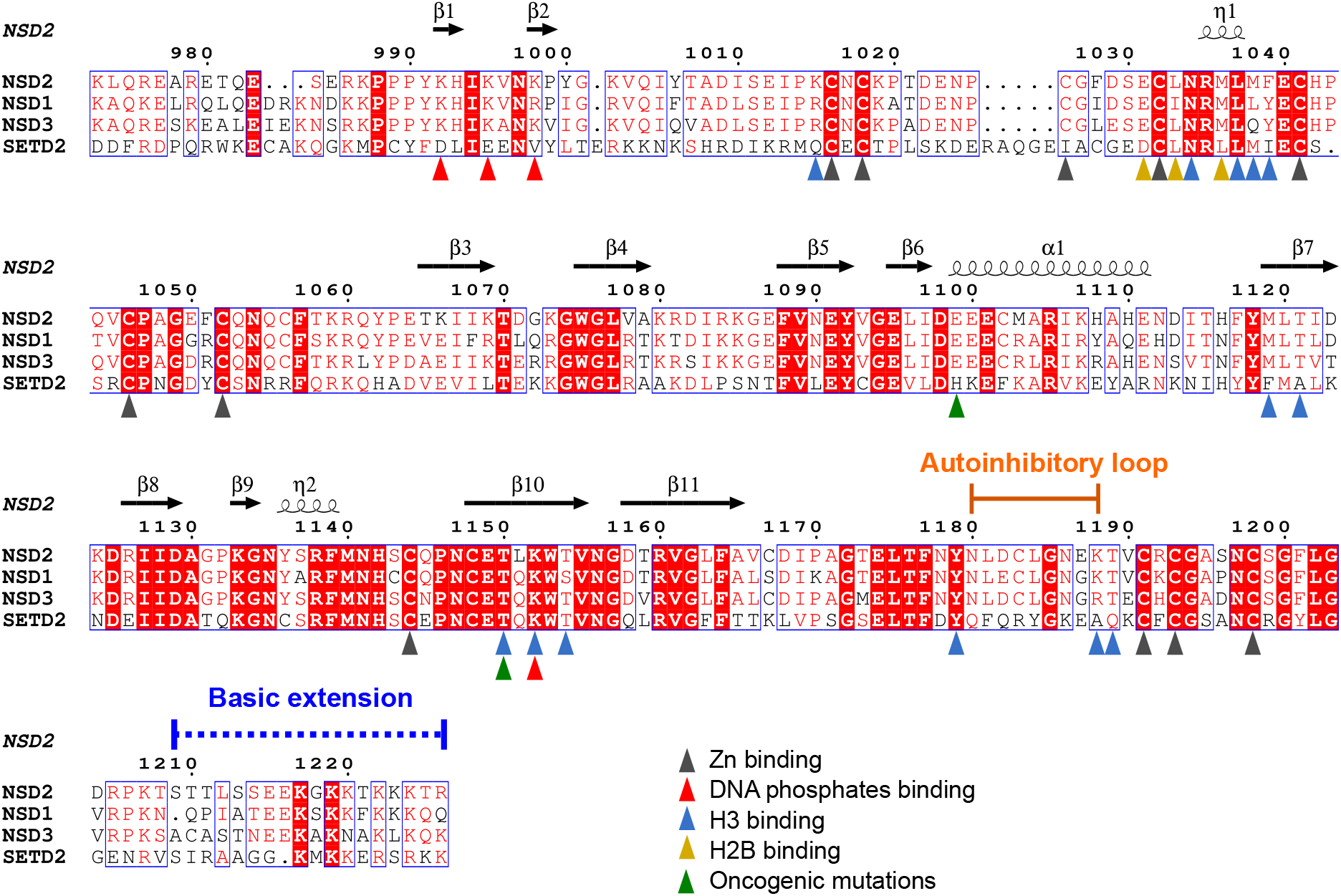
Sequence alignment of human NSD1, NSD2, NSD3, and SETD2.

**Supplementary Figure S4.**
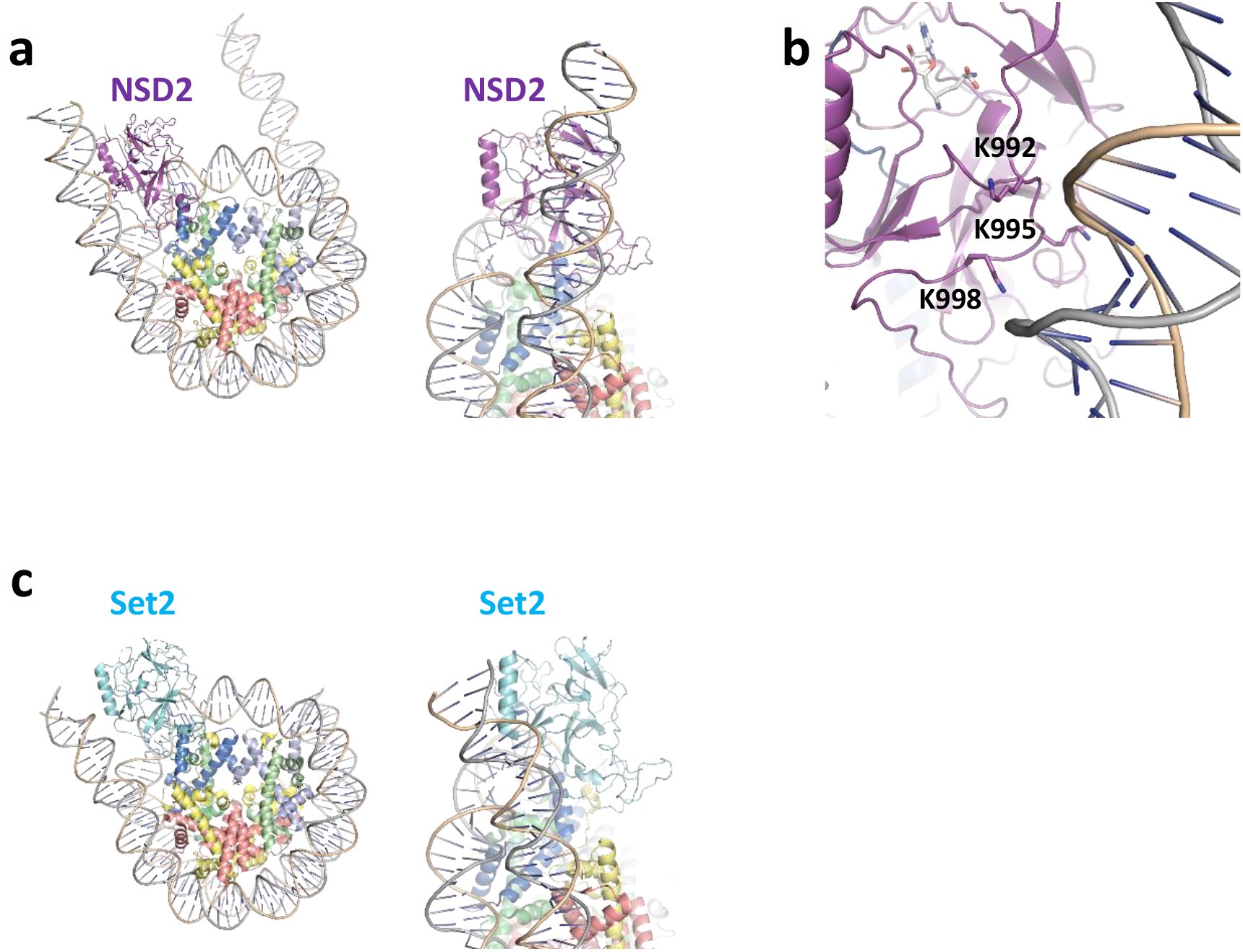
Structural comparison between NSD2 and yeast Set2. **a**, Structure of the NSD2-nucleosome complex from two views. **b**, A close-up view of the three lysine residues that interact with nucleosomal DNA in the NSD2-nucleosome complex. **c**, Structure of the Set2-nucleosome complex, same view as in (**a**).

**Supplementary Figure S5.**
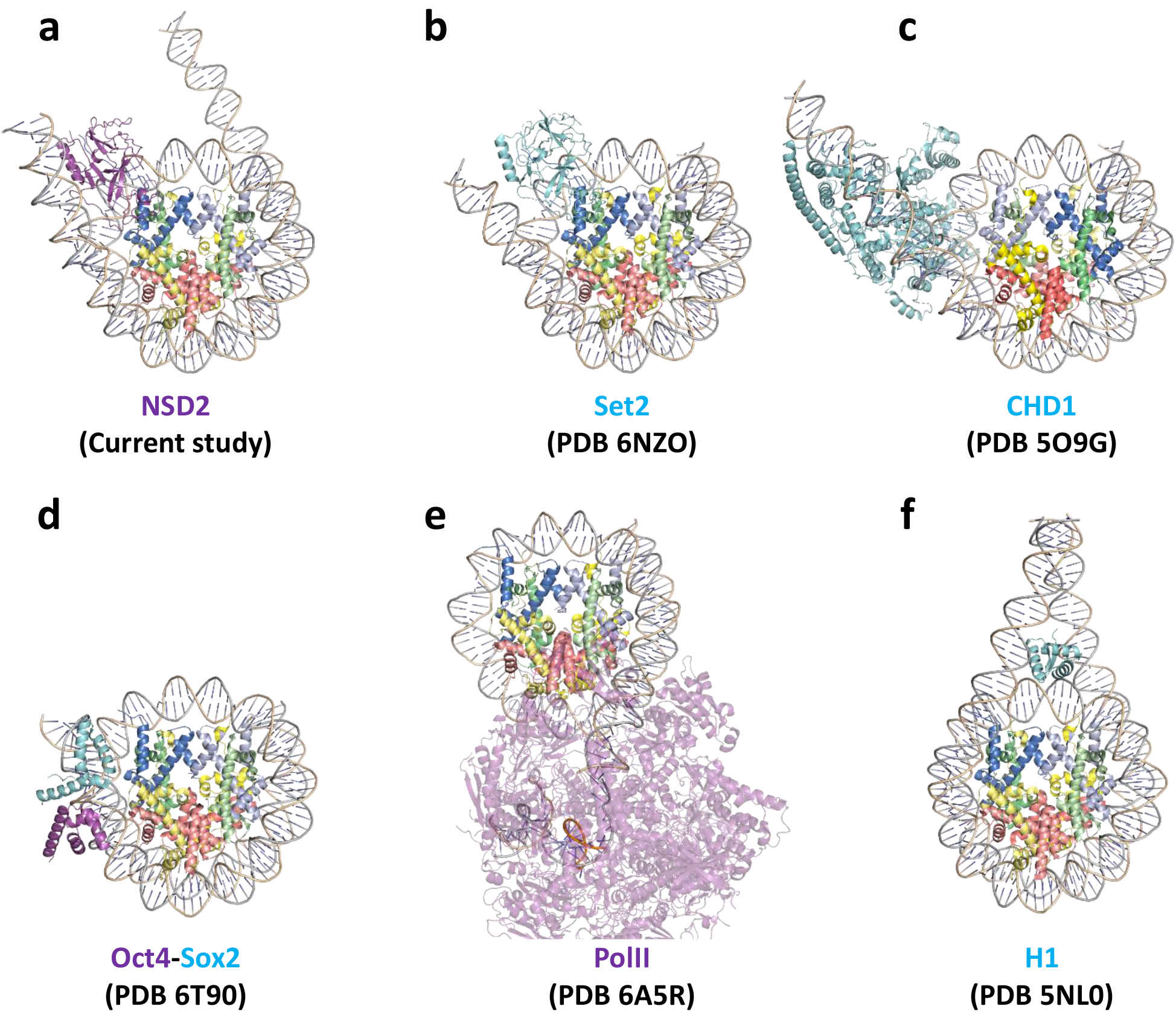
Structures of chromatin factors bound to the nucleosome. **a**, NSD2-nucleosome complex. **b**, Set2-nucleosome complex. **c**, CHD1-nucleosome complex. **d**, Oct4-Sox2 bound to nucleosome. **e**, Yeast RNA polymerase II (PolII) bound to nucleosome. **f**, Linker histone H1 bound to nucleosome.

**Supplementary Figure S6.**
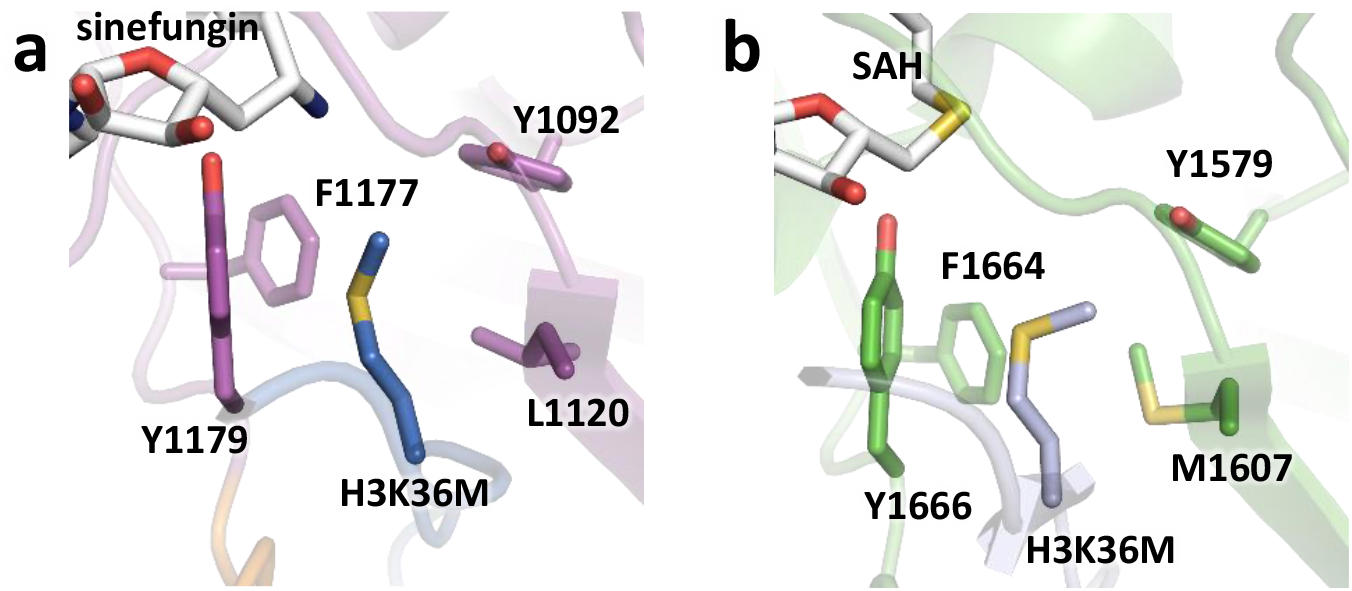
Structural comparison of the H3K36-binding cavities of NSD2 and human SETD2. **a**, NSD2 (the current structure). **b**, SETD2 (PDB ID 5JJY).

**Supplementary Figure S7.**
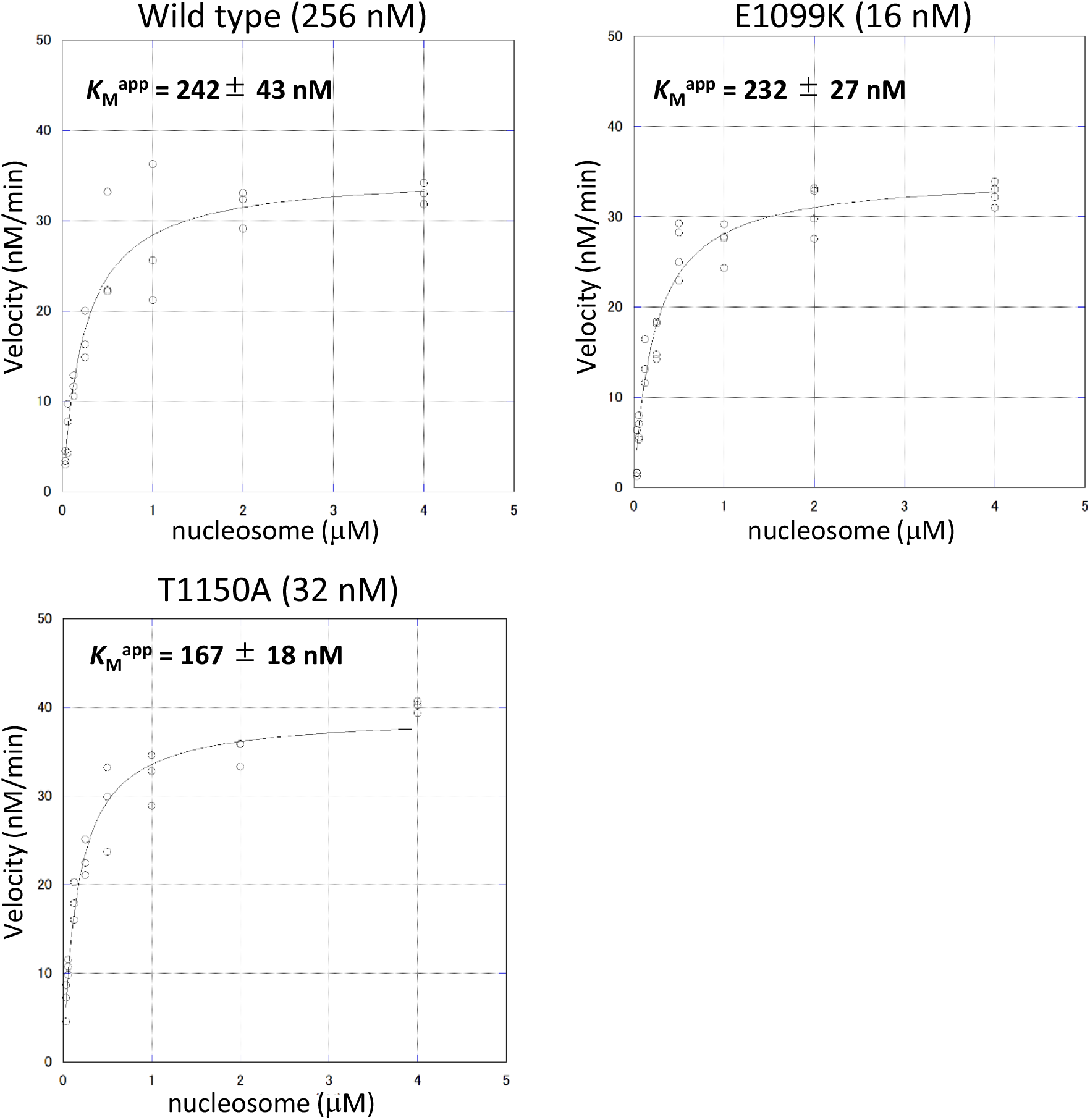
Michaelis–Menten plots using the 185-bp nucleosome as a substrate.

**Supplementary Figure S8.**
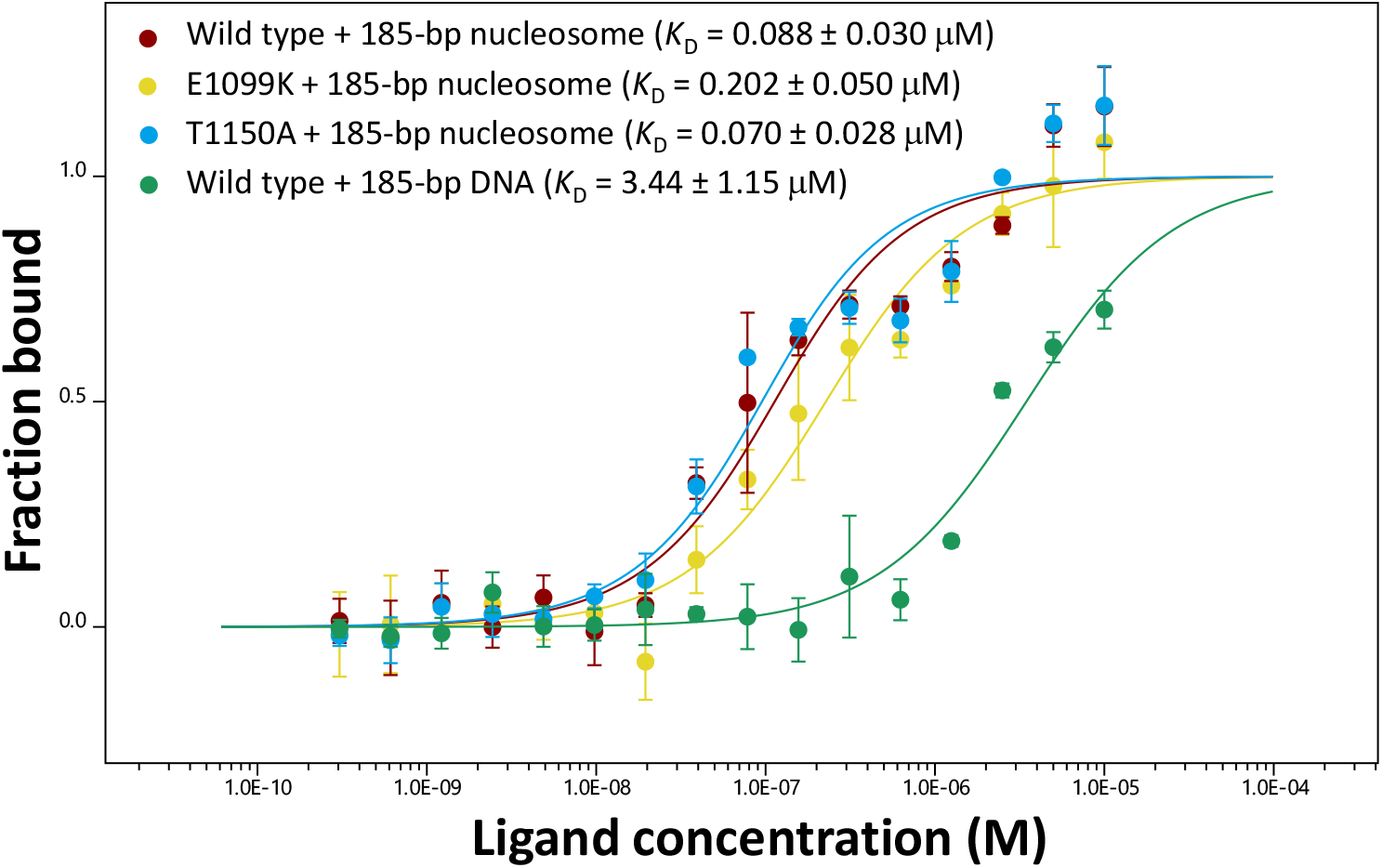
Dose-response curves of the MST assay analyzing the interactions between wild-type or mutant NSD2 and 185-bp nucleosome or naked 185-bp DNA.

**Supplementary Figure S9.**
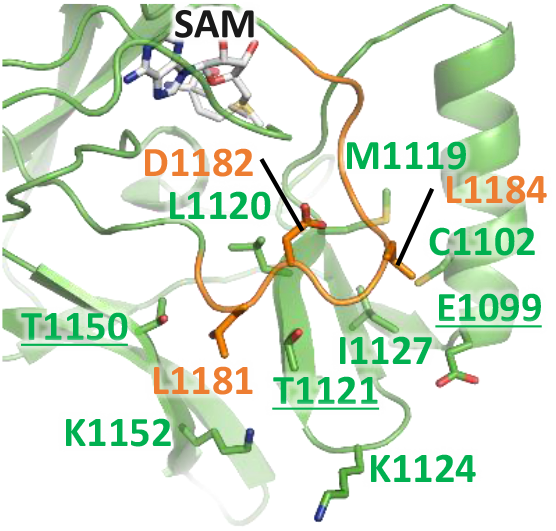
Representative structure of the wild-type NSD2 during MD simulation, in which the L1181 side chain forms hydrophobic interactions with T1121 and T1150 near the H3P38-binding patch.

**Supplementary Movie 1** Representative movie of the E1099K mutant showing autoinhibitory loop movement as it changes from a closed state to an open state, observed during the MD simulation. The carbon atoms of T1150 and E1099K are colored cyan. Backbone of the autoinhibitory loop is colored orange. Carbon atoms of the residues in H3V35-binding patch are colored yellow. L1181, C1183, and L1184 are colored according to the state of the autoinhibitory loop (red, all locks operational; white, locks partially released; blue, all locks released).

**Supplemental Table 1.** Cryo-EM data acquisition, refinement, and validation statistics

|  | 25 mM KCl (1) | 25 mM KCl (2) | 50 mM KCl |
| --- | --- | --- | --- |
| **Data collection** |  |  |  |
| Magnification | 105,000 | 105,000 | 105,000 |
| Voltage (kV) | 300 | 300 | 300 |
| Electron exposure (e<sup>-</sup>/Å<sup>2</sup>) | 72.9 | 50 | 50 |
| Frame | 70 | 48 | 48 |
| Defocus range (μm) | -0.8 to -1.6 | -0.8 to -1.6 | -0.8 to -1.6 |
| Pixel size (Å) | 0.83 | 0.83 | 0.83 |
| No. of micrographs | 2385 | 90 | 2008 |
| **Data processing** | **NSD2-nucleosome** |  |  |
| Symmetry imposed | C1 |  |  |
| Initial particle images | 4,784,875 |  |  |
| Final particle images | 136,108 |  |  |
| Map resolution (Å) | 2.8 |  |  |
| FSC threshold | 0.143 |  |  |
| **Refinement** |  |  |  |
| Initial models (PDB IDs) | 1KX5, 5LSU |  |  |
| Model composition |  |  |  |
| Non-hydrogen atoms | 14,875 |  |  |
| Protein residues | 983 |  |  |
| DNA residues | 344 |  |  |
| Ligands | sinefungin: 1<br>zinc: 3 |  |  |
| *B*-factors (Å<sup>2</sup>) |  |  |  |
| Protein atoms | 54.76 |  |  |
| DNA atoms | 139.62 |  |  |
| Ligand atoms | 101.09 |  |  |
| RMS deviations |  |  |  |
| Bonds (Å) | 0.009 |  |  |
| Angles (°) | 0.685 |  |  |
| Validation |  |  |  |
| MolProbity score | 2.31 |  |  |
| Clashscore | 8.50 |  |  |
| Ramachandran plot |  |  |  |
| Favored (%) | 93.99 |  |  |
| Allowed (%) | 5.91 |  |  |
| Outliers (%) | 0.10 |  |  |

**Supplementary Table 2.** List of MD simulation systems

| No. | Referred name | Details of the system | Duration |
| --- | --- | --- | --- |
| 1 | Wild-type | NSD2 and SAM | 500 ns × 3 |
| 2 | E1099K | NSD2 E1099K and SAM | 500 ns × 3 |
| 3 | T1150A | NSD2 T1150A and SAM | 500 ns × 3 |
| 4 | E1099K-T1150A | NSD2 with two mutations E1099K and T1150, and SAM | 500 ns × 3 |

NSD2 refers to the residues 973 to 1203 constituting N-terminal helix, AWS domain, SET domain, and the PS loop. SAM: S-adenosyl methionine. The starting structure of NSD2 was taken from cryo-EM structure reported in the current study.

**Supplementary Table 3.** Open-close dynamics of the autoinhibitory loop

| MD system | % release of the individual locks |  |  |  | All released (%) |
| --- | --- | --- | --- | --- | --- |
|  | L1184@Cγ-C1102@Sγ (D1) | C1183@Sγ-C1102@Sγ (D2) | C1183@Sγ-T1121@Cβ (D3) | L1181@Cγ-T1121@Cβ (D4) |  |
| Wild-type | 60.4 | 44.1 | 68.5 | 5.1 | 0.86 |
| E1099K | 30.3 | 81.2 | 50.5 | 74.0 | 9.7 |
| T1150A | 50.4 | 85.1 | 58.3 | 21.1 | 6.3 |
| E1099K-T1150A | 67.1 | 67.5 | 61.0 | 43.8 | 18.8 |

Note: A lock is operational when the two atoms indicated are closer than 6.5 Å; otherwise it is released. When all the locks are released, the autoinhibitory loop is considered “open”.

